# Analyzing nicotine action against amyloid toxicity by NMR-pharmacometabolomics: an exploratory study

**DOI:** 10.1101/2025.09.03.673931

**Authors:** Enza Napolitano, Carmen Marino, Manuela Grimaldi, Michela Buonocore, Anna Maria D’Ursi

## Abstract

Alzheimer’s disease (AD) is the primary neurodegenerative disease spread worldwide. One of the main histopathological hallmarks of AD is amyloid plaque deposition in the brain. Despite some epidemiological studies demonstrating that cigarette smoke is a factor in predisposing people to AD, nicotine, the principal alkaloid of *Nicotiana Tobacco*, has been widely studied for its ability to improve cognitive performance, both in animal models and in human studies.

Several hypotheses have been proposed to explain the mechanism of action underlying the beneficial effect of Nicotine in AD; however, this is still questioned.

To have new insights into the molecular mechanism underlying the neuroprotective action of Nicotine in Alzheimer’s disease, we performed an NMR metabolomic analysis of SH-SY5Y neuroblastoma cells treated with Aβ (1-42) in the presence of nicotine. Our data show that the neuroprotective action of nicotine resides in its ability to restore the systemic unbalanced metabolism associated with AD. In particular, nicotine reverses most Aβ (1-42)-induced metabolic impairments, including those related to amino acid metabolism, especially those involved in neurotransmission, as well as alterations in energy metabolism and membrane phospholipid metabolism.

## 1. Introduction

Alzheimer’s disease (AD) is the primary neurodegenerative disease spread worldwide, and it is estimated that up to 107 million subjects will be affected by 2050. ^1^

One of the main histopathological hallmarks of AD is amyloid plaque deposition in the brain, whose aggregation seems to occur decades before the disease’s onset. ^2^ Amyloid peptide (Aβ) is derived from a large protein called amyloid precursor protein (APP). In the neurons of the subjects suffering from AD, APP is cleaved firstly by the β-secretase enzyme and then by γ-secretase resulting in β-amyloid 40 and β-amyloid 42 (Aβ (1-40), Aβ (1-42)) production. ^3^ These peptides undergo a conformational transition to aggregate around meningeal and cerebral vessels and grey matter. The extracellular plaques so formed disrupt neural function, causing memory loss and cell death. ^4^

Despite several epidemiological studies demonstrating that cigarette smoke is a factor in predisposing people to neurodegenerative diseases such as AD ^5^, nicotine, the principal alkaloid of *Nicotiana Tobacco*, has been widely studied for its ability to improve cognitive performance, including attention, working memory and episodic memory both in preclinical models and in human studies. ^6^ Furthermore, nicotine treatment has been shown to reduce β-amyloid peptide (Aβ) accumulation in the cortex and hippocampus in rat models. ^7^ Published studies in humans have reported that intravenous and subcutaneous nicotine administration in AD patients improved several cognitive tasks, such as visual attention and perception ^8^ and mood and lexical tasks ^9^ although not memory. ^10^

The mechanism of action responsible for the beneficial effect of nicotine in AD preclinical models and patients is still questioned, although some hypotheses have been proposed: i) the concentration of nicotinic ACh receptors (nAChRs) are selectively decreased in the AD brain^11^ so that the beneficial effects of nicotine could depend on an up-regulation of nicotine receptors ^12^; ii) nicotine decreases the accumulation of Aβ in the cortex and hippocampus of mice models of AD, preventing the activation of NF-κB and c-*Myc* by inhibiting the activation of MAP kinases (MAPKs). Thus, inducible NOS and NO production activity are downregulated. ^13^ iii) Furthermore, nicotine binding to a7nAChR prevents Aβ interaction with nicotine receptors, which causes the inhibition of a7nAChR-dependent calcium activation and the acetylcholine release, two processes critically involved in memory and cognitive functions. ^14^ iv) enhanced oxidative stress characterized the brain of AD patients, and some studies suggest that the beneficial effects of nicotine in neurodegenerative disease may be, at least partly, due to an antioxidant mechanism. ^15^; v) the neuroprotective effect of nicotine resides in its antiaggregant properties. Structural studies investigating the interaction of Aβ peptides with nicotine and its derivatives demonstrate that nicotine slows down the aggregation of Aβ (1-42) ^16^ and Aβ (25–35) peptides ^17^.

Recently, pharmacometabolomic has emerged as an innovative approach for analyzing the molecular mechanism and toxicity of approved drugs and new molecular entities ^18^. Beside the applications in structural analysis, NMR spectroscopy represents a robust and suitable technique for metabolomic studies, enabling the simultaneous qualitative and quantitative identification of low-molecular-mass compounds in biofluids and other biological samples. ^19^

To have new insights into the molecular mechanism underlying the neuroprotective action of Nicotine in Alzheimer’s disease, we performed an NMR metabolomic analysis of SH-SY5Y neuroblastoma cells treated with Aβ (1-42) in the presence of nicotine. SH-SY5Y cells are often used as a cellular model for studying Alzheimer’s disease (AD) and other neurodegenerative disorders. Our data confirm the results of previous metabolomic studies performed on cell cultures exposed to Aβ (1-42) and on biological fluids of animals and humans ^20–22^. Interestingly, our data prove a rebalancing of the metabolic condition of SH- SY5Y cells pretreated with nicotine and incubated with Aβ (1-42) towards the metabolic condition of the healthy control cells. A careful analysis of our data to understand how nicotine may impact different sides of cellular metabolism suggests significant effects of the alkaloid on (i) amino acid metabolism, particularly those involved with neurotransmission, (ii) energy metabolism, and (iii) membrane phospholipid metabolism.

## 2. Materials and Methods

### 2.1 Chemicals

Dulbecc’s Modified Eagle’s Medium (DMEM), glutamine, penicillin and streptomycin, fetal bovine serum (FBS), CCK-8, (-)-Nicotine (≥99%) were purchase from Sigma-Aldrich (St. Louis, MI, USA). Aβ (1-42) peptide was expressed as previously reported.^23^

### 2.2 Cell Culture and Drug Treatment

Undifferentiated SH-SY5Y human neuroblastoma cells were obtained from American Type Culture Collection (ATCC, Rockville, MD, USA). Cells were cultured in Dulbecco’s Modified Eagle Medium (DMEM, 4500 mg/mL glucose) supplemented with 10% (*v*/*v*) fetal bovine serum (FBS), 2 mM L-glutamine, 100 U/mL penicillin, and 0.1 mg/mL streptomycin. Cells were grown in 100 mm culture dishes and incubated at 37 °C under a humidified atmosphere with 5% CO2, until reaching semiconfluency and split every 2 days.

### 2.3 Cell Viability Assay

Cell viability was established by measuring mitochondrial metabolic activity with Cell Counting Kit-8 (CCK-8 Cat. CK04, Dojindo Laboratories, Rockville, MD, USA). ^24^ This assay evaluates the viability of cells considering the ability of dehydrogenases’ cells to reduce the tetrazolium salt WST-8 (2-(2-methoxy-4-nitrophenyl)-3-(4-nitrophenyl)-5-(2,4- disulfophenyl)-2H-tetrazolium, monosodic salt) in an orange-colored formazan dye, which is soluble in the tissue culture medium. The quantity of formazan dye generated by intracellular dehydrogenase activity is directly proportional to the number of living cells.

Briefly, SH-SY5Y (8 × 10^3^ cells/well) were plated into 96-well plates for 24 h, then nicotine 1 mM or 100 μM was added for 24 h. Next, Aβ (1-42) peptide 25 μM was added for 48 h. At the end of this period, CCK-8 reagent at 10% final concentration was added for 1 h. Afterward, the absorbance was measured at 450 nm, using a microplate reader (Multiskan Go, Thermo Scientific, Waltham, MA, USA). Cell viability was expressed as a percentage relative to the untreated cells cultured in medium only with vehicle and set to 100%, whereas 10% DMSO was used as positive control and set to 0% of viability. Results are expressed as means ±SD of 3 independent experiments performed in triplicate and reported as percentage of viable cells vs the untreated control.

### 2.4 Exposure of SH-SY5Y to Nicotine and Aβ (1-42)

Cells were pleated in 60 mm culture dishes and allowed to adhere overnight. For the nicotine Aβ (1-42) co-administration, cells were pretreated with nicotine (100 μM) and after 24 h Aβ (1-42) was added for additional 48 hours at a sub-toxic concentration (5 μM). Cells exposed only to Aβ (1-42) peptide at the same concentration and incubation time were used for the comparison. For the control group cells were treated only with vehicle. At the end of treatments, the medium was collected, and the dishes washed with cold PBS (pH 7.4) to remove media components, immediately before the solvent extraction procedure described below.

All conditions were tested in three biological replicates, and each biological replicate provided three technical replicates.

### 2.5 Sample extraction

The culture medium was collected from each plate (including cell-free medium incubated under the same conditions) in microcentrifuge tubes and centrifuged at 1000× g for 10 min. For extracting cellular metabolites, a biphasic extraction protocol (methanol:chloroform:water; 1:1:1) was used, after cell collection by scraping and homogenization (Beckonert et al., 2007). After centrifugation at 6000 rpm for 10 min at 4 °C the two phases were separated. The resulting polar extracts were dried under vacuum in a SP- Genevac EZ-2 4.0 concentrator, and the lipophilic extracts were dried under a nitrogen flow for future analysis. All extracts were stored at −80 °C prior to NMR analysis.

### 2.6 NMR spectroscopy data acquisition and processing

Lyophilized cell extracts were dissolved in 200 μL of buffer (50 mM Na_2_HPO_4_, 1 mM trimethylsilyl propionic-2,2,3,3-d_4_ acid, sodium salt (TSP-d_4_), 10% of D_2_O) and transferred into 3 mm NMR tubes for ^1^H NMR detection. TSP-d_4_ at 0.1% in D_2_O was used as an internal reference for the alignment and quantification of NMR signals. For the extracellular analysis, 100 μL of cell medium was mixed with 100 μL of the same buffer used for the lyophilized extracts. 1D ^1^H NMR experiments were acquired using a Bruker Ascend™ 600 MHz spectrometer with a 5 mm triple resonance Z gradient TXI probe (Bruker Co, Rheinstetten, Germany) at 298 K.

Spectra acquisition was performed using 12 ppm spectral width, 20k data points, presaturation during relaxation delay and mixing time for water suppression ^25^ and spoil gradient, 5 s relaxation delay, and mixing time of 10 ms.

Topspin version 3.0 (Bruker Biospin) was used for spectrometer control and data processing. Analysis of the NMR spectra was performed based on a targeted metabolomic approach. Accordingly, each metabolite was identified before statistical analysis using Chenomx NMR-Suite v8.0 software (Chenomx NMR suite, v8.0, Edmonton, AB, Canada) combining advanced analysis tools with a compound library. Quantitative analysis of NMR spectra was performed using NMRProcFlow ^26^ and obtained data matrix was subjected to statistical analysis.

### 2.7 Statistical analysis

Sample data were normalised using sum, Log transformed, and Pareto scaled and analysed by the open-source tool Metaboanalyst 6.0 and MixOmics R-package (mixOmics-package). ^27, 28^ To increase the accuracy and biological understanding of the data, multivariate statistical analysis was first performed on exometabolome and endometabolome concentration matrices and then on combined data matrices.

Univariate analysis was conducted separately on the exometabolome and endometabolome of the groups analyzed by T-test and Fold Change and plotted by Volcano plot. ^29^

To provide an exhaustive representation of the quantitative variations of the metabolites, we carried out heatmaps using normalized data, average group concentration, and Euclidian distance. ^30^

Multivariate analysis was performed on combined matrices of endo and exometabolome using the supervised Sparse Partial Least Squares (or Projection to Latent Space-sPLS) approach. ^31^ This approach represents a linear and multivariate visualisation technique for integrable datasets that improves the shortcomings of Principal Component Analysis methods and Canonical correspondence analysis (CCA) ^32^ In the integrated approach, sPLS analysis performs well when the sum of variables in the integrated matrices exceeds the number of samples as analysed in this study. sPLS was performed considering the LASSO penalty on loading vectors to reduce the number of original variables used to construct latent variables.33

A sample plot highlighted the clustering of samples’ metabolomic profiles. In the graph, samples are represented as points positioned based on their projection of the selected latent components of the data. Leave-one-out cross-validation is performed to validate the model according to R2, Q2, and accuracy values ^34^. Furthermore, Spls models were confirmed according to distance matrices calculated by the centroid method, maximum distance, and Mahalanobis distance (**Supplementary Figure 1**). ^35^

Variable correlations were represented using a circular correlation plot. All resulting vectors are plotted within a unit circle (radius of 1). The position of each vector corresponds to its correlation with the components. Therefore, stronger associations will give rise to a vector that extends further from the origin. Moreover, variable vectors close to each other will be highly correlated. ^27^

The contribution of each variable is reported in a bar graph. The contribution graph of the variables constructed on the loadings of separation of the variables has been coloured considering the maximum against two, therefore representing the clusters in which the metabolite has the highest concentration. For 3 clusters, both labelling methods were used; the maximum and minimum concentration of the discriminating metabolite was found.

Ggalluvionalplot, a package of ggplot2 in R ^36^, has been used to graph the variations of the discriminating metabolites that have been classified considering the loading parameter resulting from the sPLS-DA and the cell compartment where down or upregulation is observed.

The enrichment pathway tool was applied to perform pathway analysis using Metaboanalyst 6.0. KEGG paths were chosen according to the lower false discoveries (FDR), p-value <0.05 and the hits value, related to the number of metabolites belonging to the pathway, >1 (**Supplementary Table 1**). ^37^

## 3. Results

### 3.1 Nicotine protects neuroblastoma cells from Aβ (1-42) toxicity

The SH-SY5Y neuroblastoma cell line (ATCC, Rockville, MD, USA) is a valuable, widely used model for studying neurodegenerative diseases like Parkinson’s and Alzheimer’s at a cellular level. ^38^

As a preliminary step, the toxicity of Aβ (1-42) recombinant protein was evaluated on SH- SY5Y cells, showing an EC_50_ of 46.75 ± 4.01 μM. (**Supplementary Figure S2**)

A viability assay was performed to estimate the protective effect of nicotine on SH-SY5Y neuroblastoma cells. We confirmed the protective effect of Nicotine against Aβ (1-42) toxicity, as reported in the literature ^39^. **Figure 1** shows that SH-SY5Y neuroblastoma cells treated with Aβ (1-42) (25 μM) have 70.12 ± 3.07% survival. The presence of 1mM and 100µM Nicotine preserved cell viability up to 90.75 ± 3.14% and 93.75 ± 1.22%, respectively.

**Figure 1.**
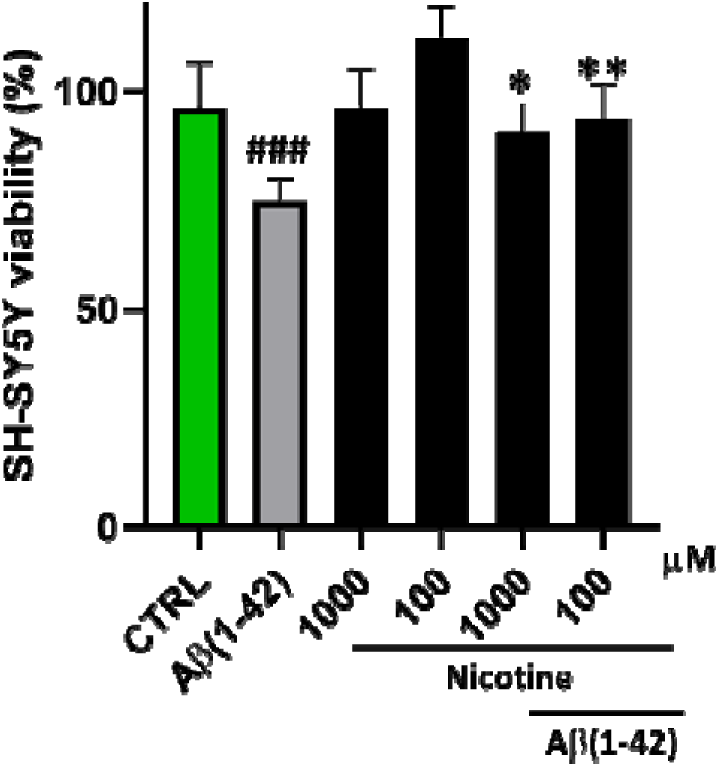
Neuroprotective effect of the Nicotine against Aβ (1-42)-induced cytotoxicity. Cell viability was examined by the CCK-8 assay. SH-SY5Y cells were exposed to nicotine (1 mM and 100μM) for 24 h before administration of Aβ (1-42) recombinant monomer 25 μM for an additional 48 h. The viability variations were calculated as the percentage of viable cells in treated cultures compared to untreated ones (CTRL). Results are shown as mean ± standard deviation (SD) from three independent experiments. ### denote respectively *p* < 0.001 vs. Ctrl; *, ** denote respectively *p* < 0.05 and *p* < 0.01 vs Aβ (1-42).

### 3.2 Aβ (1-42) disrupts energetic pathways, amino acid metabolism, and membrane stability

To evaluate the effects of Aβ (1-42) on SH-SY5Y cells, we analyzed the metabolic profile of SH-SY5Y cells exposed to Aβ (1-42) in comparison to the metabolic profile of the control cells treated with the vehicle alone (CTRL). ^1^H NMR spectra of intracellular extracts (endometabolome) and extracellular media (exometabolome) were collected (**Figures 2A, B**), and the ^1^H resonance assignment resulted in metabolite matrix including for each sample, the intracellular and extracellular metabolite concentrations.

**Figure 2.**
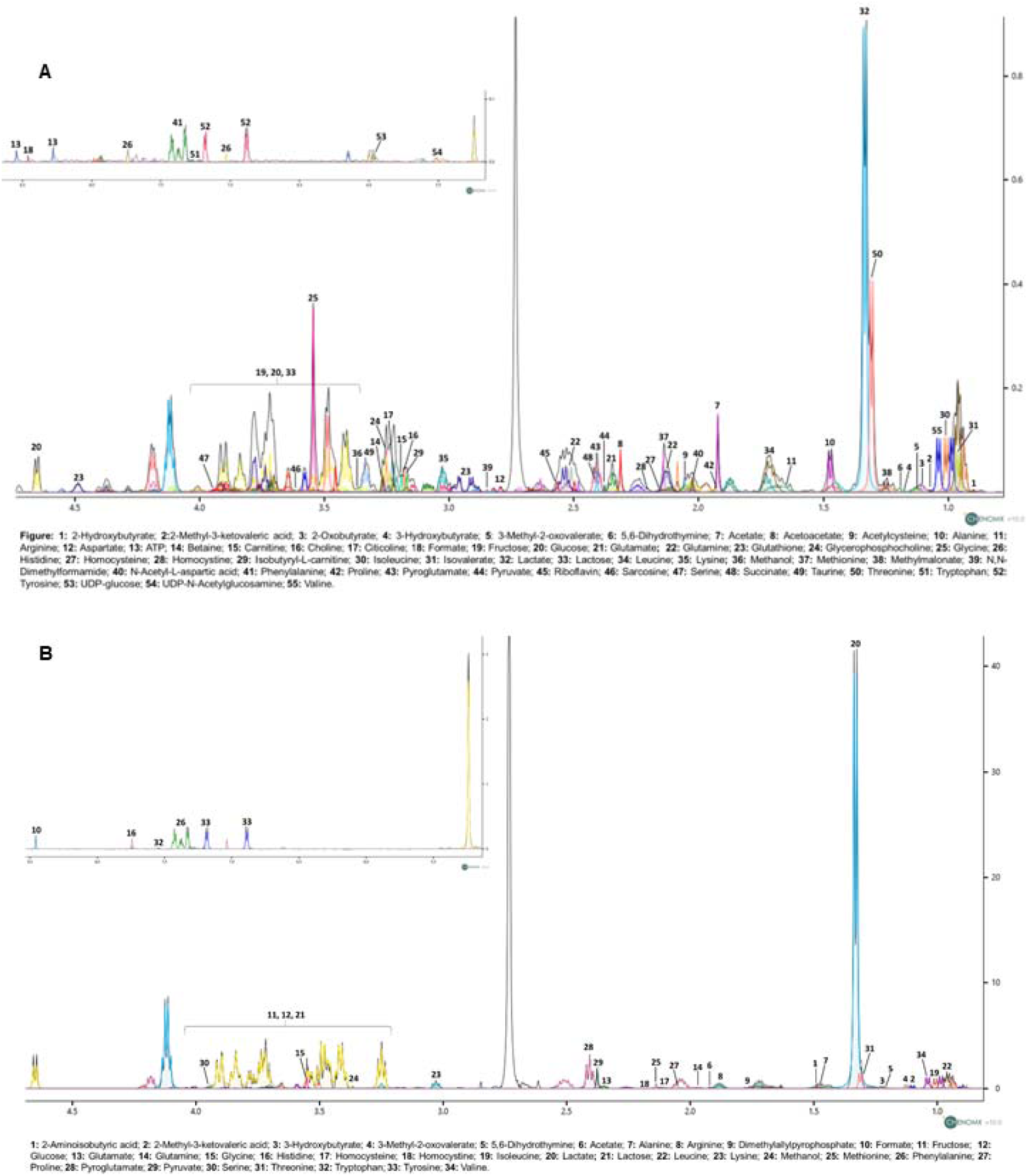
Representative 1D ^1^H NOESY spectra of polar cellular extracts of SH-SY5Y cells (**A**: endometabolome) and conditioned growth medium (**B**: exometabolome). The spectra are acquired at 600 MHz and T = 310 K. Numbers indicate metabolites listed below the spectra.

**Figures 3A** and **3B** show the Volcano Plots related to the endo and the exometabolome of cells incubated with Aβ (1-42). The quantification of the metabolite concentration reveals an increase in Pyruvate, Succinate, Isobutirril-Carnitine, and Homocysteine and a reduction of Pyroglutamate, Taurine, and Carnitine in the endometabolome. Moreover, an increase in Threonine and a decrease in Fructose in the exometabolome was detected.

**Figure 3.**
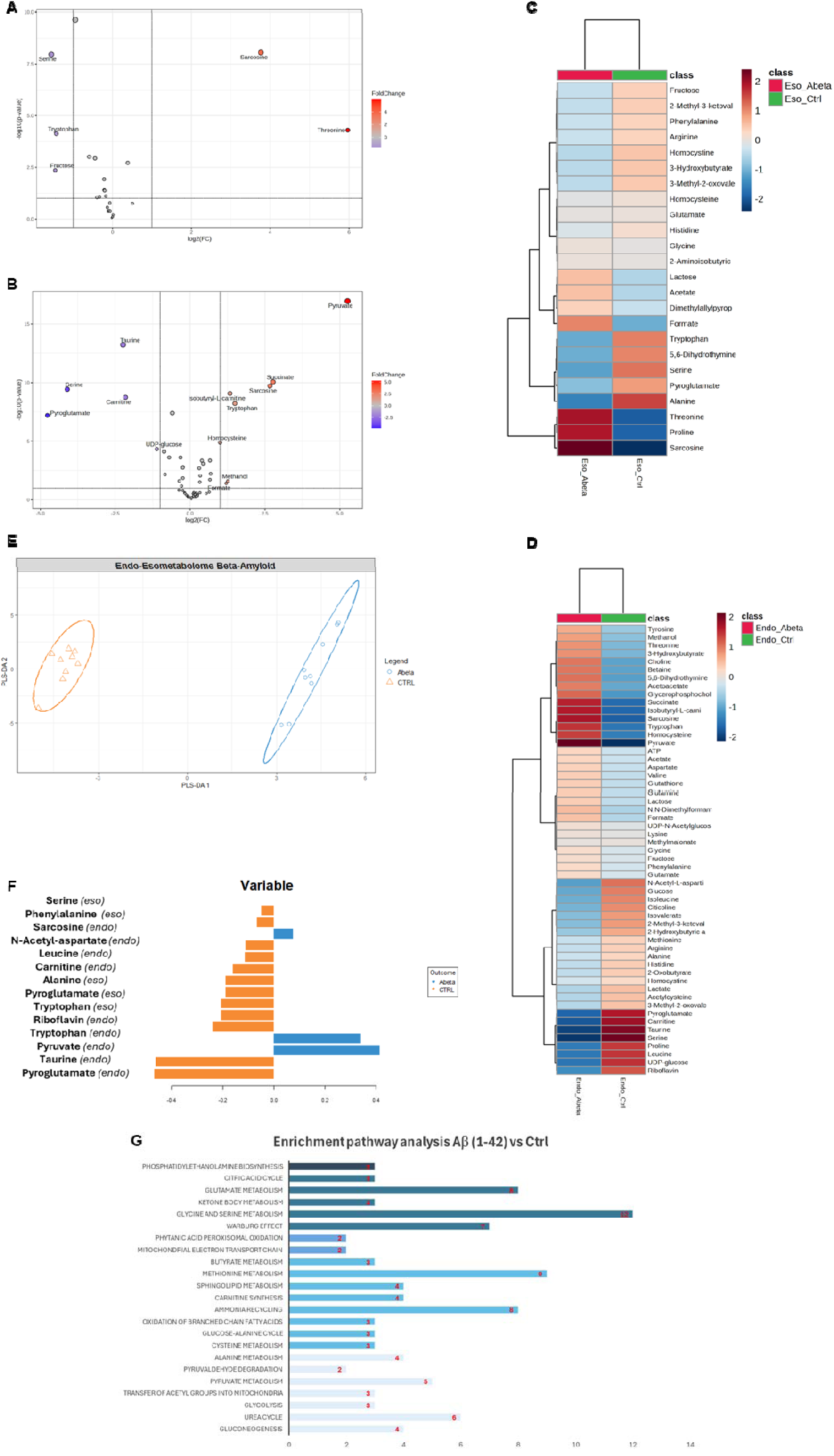
**A,B**; Volcano plot analysis of metabolic changes in the exo and endometabolome of SH-SY5Y cells incubated with Aβ (1-42) vs. control cells (Ctrl). Each point on the volcano plot was based on p-value and fold-change values, set at 0.05 and 2.0, respectively. Red points identify upregulated metabolites, whereas blue points identify down-regulated metabolites. **C,D** Heatmap of changed metabolites in exo and endometabolome respectively. The colour of each section corresponds to a concentration value of each metabolite calculated by a normalized concentration matrix (red, upregulated; blue, downregulated) **E** sPLS-DA score scatter plots related to the combined matrices of endo and exometabolome of SH-SY5Y cells treated with Aβ (1-42) (Abeta) vs. control cells (CTRL). The cluster analyses are reported in the Cartesian space described by the principal components PC1:40% and PC2:16%. sPLS-DA was evaluated using cross-validation (CV) analysis. CV tests performed according to the sPLS-DA statistical protocol show a significant cluster separation (1.0 accuracy values on PC1 and PC2, with positive 0.80 and 0.90 Q2 indices, respectively). **F** Loadings barplot related to the combined matrices of endo and exometabolome. The variables responsible for metabolomic profile differences are ordered according to values of increasing importance from bottom to top. Colors indicate the cluster where the median is maximum for each metabolite (blue: Abeta; orange: CTRL). **G** Enrichment pathways analysis: the discriminative pathways are ranked according to p.value and number of hits reported in the bars.

Based on univariate statistical analysis, sarcosine appears to increase and serine to decrease in both endo and exo metabolomes of the SH-SY5Y cells incubated with Aβ (1-42); conversely, tryptophan concentration increases in the intracellular compartment and decreases in extracellular one.

Volcano plot’s results were confirmed by heatmap analysis, indicating, with a specific color code, the downregulation (in blue) and up regulation (in red) of all the metabolites detected in the spectra. (**Figures 3C, D**)

Using a combined approach in analyzing the metabolomic profile of intra- and extracellular compartments, we derived the sPLS-DA score plot representing the metabolomic profile of the cell compartments of SH-SY5Y cultures incubated with Aβ (1-42) vs. untreated control (**Figure 3E**). The Cartesian space is described by the first and second main components (PC1; PC2) and explains 40% and 16% of dataset variance. The model’s validity was evaluated through a cross-validation approach using Mahalanobis distance, maximum distance, and centroids (**Figure S1**). The separation of clusters indicates that Aβ (1-42) perturbs both exometabolome and endometabolome of the cell cultures.

The variables’ loadings were calculated to identify the metabolites responsible for cluster separation. The bar plot, shown in **Figure 3F**, reports the discriminating metabolites classified by their loading values and the clusters where the concentration of each metabolite is highest.

The data indicated a reduction in the concentrations of Taurine, Riboflavin, Carnitine, Leucine, and N-acetylAspartate (NAA) in the endo metabolome of cells treated with Aβ (1- 42); on the contrary, an increase in the Pyruvate, and Sarcosine concentrations was observed respect with control cells.

Furthermore, the exometabolome of cells incubated with Aβ (1-42) has lower Alanine and Phenylalanine concentrations than untreated cells. The combined analysis also showed an influence of Aβ (1-42) in reducing the concentrations of Pyroglutamate and Serine in both cellular compartments. In contrast, Tryptophan concentration is reduced in exometabolome and increased in endometabolome. (**Figura 3F**)

Pathways analysis carried out on the endometabolome’s quantified metabolites showed an impact of Aβ (1-42) on the membrane’s lipids metabolism in *Phosphatidylethanolamine biosynthesis and sphingolipid metabolism.* Enrichment Pathways also points to energetic metabolic dysregulation mainly linked to the *Citric Acid Cycle, Ketone Body metabolism, Pyruvate metabolism, Glycolysis* and *Urea cycle*. In addition, dysregulation of several amino acid pathways, such as *Glutamate Metabolism, Glycine and Serine metabolism,* and *Methionine metabolism*, has also been reported. (**Figure 3G**)

### 3.3 Nicotine reverts AD dysmetabolism, acting on the whole metabolome

To evaluate the potential of nicotine in modulating the dysmetabolism caused by Aβ (1-42), we treated SH-SY5Y with nicotine for 24 hours before exposure to Aβ (1-42). The metabolite concentrations of intra and extracellular compartments, derived from quantitative analysis of ^1^H NMR spectra, were analyzed using multivariate statistical analysis (MVA). To examine the impact of nicotine on Aβ (1-42)-induced alterations on SH-SY5Y metabolism, we conducted a comparative analysis of the endo and exometabolome profiles of three distinct cell groups: (i) cells treated with nicotine and then incubated with Aβ (1-42) (Abeta_Nic); (ii) cells incubated with Aβ (1-42) alone (Abeta); and (iii) untreated cells (CTRL).

The combined omic analysis using sPLS-DA shows a clear separation of the three metabolomic profiles. This indicates that the effect of nicotine on cells incubated with Aβ (1- 42) is different from control cells, which represent the healthy cellular phenotype. (**Figure 4A**) The model’s validity was estimated through a cross-validation approach using Mahalanobis distance, maximum distance, and centroids (**Figure S2**).

**Figure 4.**
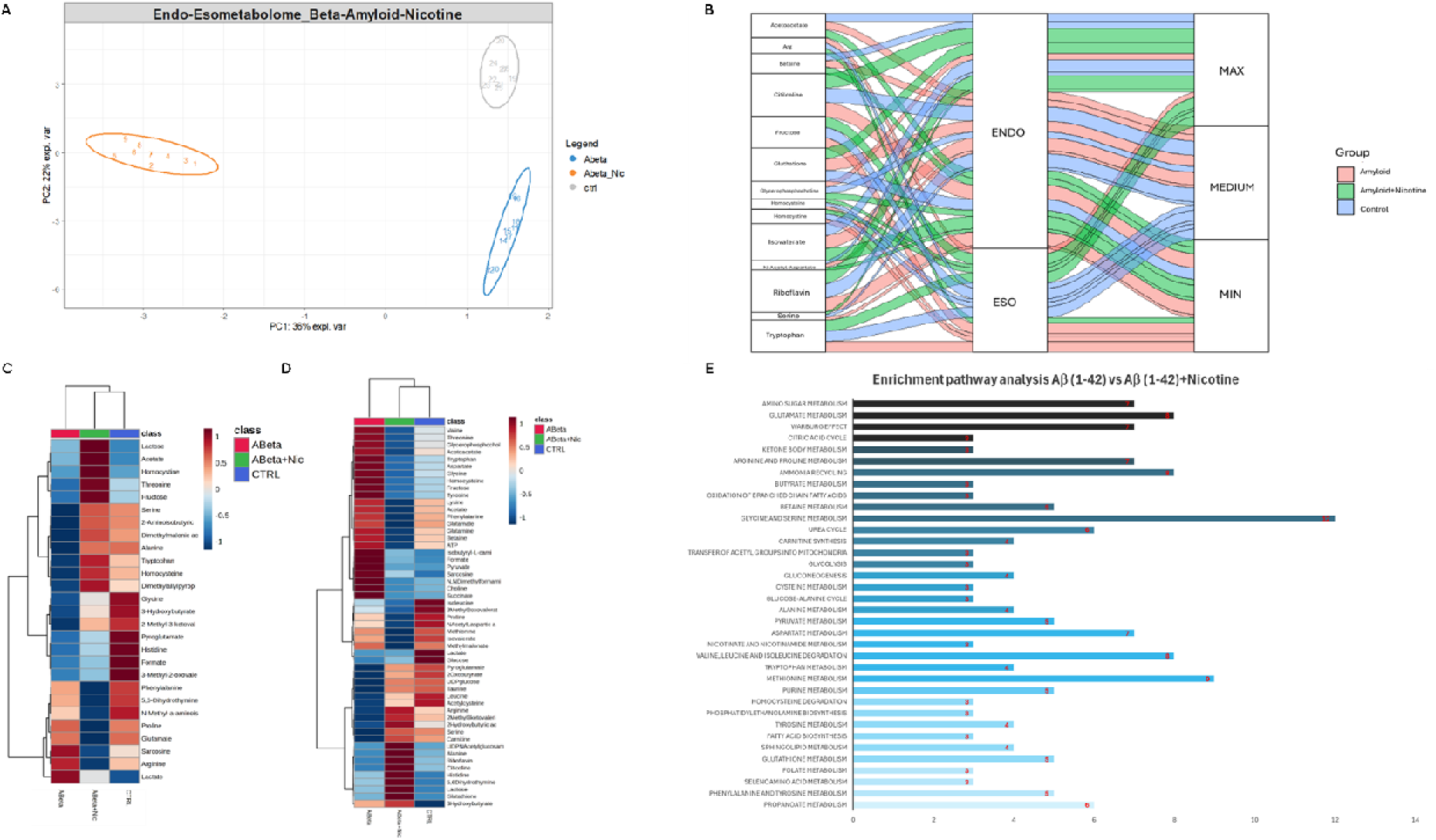
**A** sPLS-DA score scatter plots related to the combined matrices of endo and exometabolome of SH- SY5Y cells treated with Aβ (1-42) in blue (Abeta) vs. cells pretreated with nicotine before being incubated with Aβ (1-42) in orange (Abeta_Nic) vs. control cells in grey (CTRL). The cluster analyses are reported in the Cartesian space described by the principal components PC1:36% and PC2:22%. sPLS-DA was evaluated using cross-validation (CV) analysis. CV tests performed according to the sPLS-DA statistical protocol show a significant cluster separation (0.60 and 1.0 accuracy values on PC1 and PC2, with positive 0.77 and 0.95 Q2 indices, respectively). **B** Alluvional plot reporting metabolites discriminating clusters analyzed in sPLS-DA and classified according to loading value. The first column reports the discriminating metabolites, the second column is the cellular compartment, and the third column is the concentration change. The maximum concentrations in the comparison between the three clusters are indicated with "Max", the intermediate with "Medium", and the minimum with "Min". The lines connect the metabolite, the cell compartment where it is discriminating and the column representing quantitative variation. Each line is reported in pink if the metabolite has the variation in the clusters of cells incubated with Aβ (1-42); in green if the variation is typical of cells pretreated with nicotine before being incubated with Aβ (1-42) and in blue if the variation is typical of control cells **C, D** Heatmap of changed metabolites in exo and endometabolome respectively. The color of each section corresponds to a concentration value of each metabolite calculated by a normalized concentration matrix (red, upregulated; blue, downregulated **E** Enrichment pathways analysis: the discriminative pathways are ranked according to p.value and number of hits reported in the bars

The alluvial plot shown in **Figure 4B** reveals that the endometabolome of the cells pretreated with nicotine before being incubated with Aβ (1-42) have high concentrations of Citilcoline, Glutathione, Riboflavin, and Serine. In contrast, the endometabolome of this group of cells includes a minimal concentration of Acetoacetate, Betaine, Glycerophoshocholine, Isovalerate, and N-acetylaspartate (**Figure 4B**). Furthermore, the exometabolome of SH- SY5Y cells incubated with nicotine and Aβ (1-42) has the highest concentration of Fructose, Homocysteine, and Tryptophan, while low concentration of Arginine compared to the same cells incubated with Aβ (1-42) in the absence of nicotine or compared to the untreated cells.

Heatmaps (**Figures 4C, D**) based on average concentrations of the cytoplasmic and extracellular metabolite concentrations of the three groups of cells under investigation show that the metabolomic profile of the cells treated with Aβ (1-42) in the presence of nicotine is very similar to the control cells.

As shown in **Figures 4C** and **4D**, the exometabolome and endometabolome of neuroblastoma cells treated with nicotine and Aβ (1-42) cluster with the control cells’ metabolomic profile. This suggests that nicotine induces metabolic changes towards a phenotype more similar to healthy control rather than the pathological AD condition, represented by cells exposed only to Aβ (1-42).

**Figure 4E** reports the results of the Enrichment pathway analysis performed on the data matrix containing the endometabolites relative to the cells i) pretreated with nicotine before being incubated with Aβ (1-42) ii) incubated only with Aβ (1-42) to evaluate the biochemical pathways regulated by nicotine in the AD cellular model.

Accordingly, it is evident that there is a dysregulation of *Betaine metabolism*, *Methionine metabolism, Homocysteine degradation,* and *Folate metabolism.* Moreover, an alteration of several energetic pathways, such as *Amino sugar metabolism, Citric acid cycle, Ketone body metabolism, Glycolysis*, and *Pyruvate metabolism* has been shown.

Enrichment analysis revealed a modulation of the amino acid pathways most involved in neurotransmission, such as *Glutamate metabolism* and *Glycine and serine metabolism*, as well as *Phenylalanine and tyrosine metabolism*. Furthermore, nicotine has been demonstrated to exert an influence on energetic pathways, including *the Citric acid cycle, Ketone body metabolism* and *Urea cycle*. Moreover, the Enrichment analysis confirmed the effect on the pathways of membrane phospholipid and sphingolipid biosynthesis and revealed an antioxidant action of nicotine, identifying the dysregulation of *Glutathione metabolism*.

## Discussion

The pyridine alkaloid nicotine has been blamed for a long time for its association with smoking and addiction. Recently, a renewed interest emerged in the investigation of Nicotine as a compound endowed with biological activity in controlling AD symptoms.

In particular, nicotine appears to have an impact in controlling neuroinflammation and apoptosis and reducing the misfolding of amyloid proteins. ^7, 13, 16, 17, 40^ SH-SY5Y cells are recognized as a functional cellular model for studying neurogenerative diseases like AD and Parkison’s. ^38^ This cell line has proven to be a valuable model for studying the metabolic changes associated with Alzheimer’s disease (AD) when treated with the amyloid peptide Aβ (1-42) ^20, 21^

To gain insights into the hitherto unknown role of nicotine in protecting from the neurotoxic action of Aβ (1-42) amyloid peptide, we performed an NMR-based metabolomics investigation of SH-SY5Y neuroblastoma cells exposed to nicotine before being treated with Aβ (1-42).

In recent years, metabolomics has emerged as a valuable method for investigating the molecular mechanism of active biological compounds. Indeed, an innovative aspect of metabolomics lies in its untargeted nature, potentially allowing the disclosure of unexplored connections between disease, therapeutics, and biological pathways.

Data previously acquired using metabolomic analysis of SH-SY5Y cell treated Aβ (1-42) resemble AD alterations reported in patients’ biofluids, such as increased oxidative stress and inflammation, with unfavorable effects on neurotransmission and lipid metabolism. ^21, 41^ This effect is particularly evident in altered phosphatidylcholine and lipo-phosphatidylcholine concentrations and coincides with data also derived from CSF and plasma analysis of AD patients. ^20^

Our NMR-based metabolomic investigation was exhaustive and took advantage of a combined approach focused on the endo- and exometabolomic of SH-SY5Y cells treated Aβ (1-42). In agreement with the evidence previously collected from experiments on AD cellular models and Alzheimer’s patient biofluids, our observations pointed to abnormally low NAA levels that were compatible with altered glutamatergic neurotransmission. ^21, 42^ Changes in NAA could impact glutamate neurotransmission, as NAA may function as a neurotransmitter in the brain by modulating metabotropic glutamate receptors. ^43^ Furthermore, we observed that Aβ (1-42) induces increased sarcosine and decreased Serine and Pyroglutamate concentrations in both extra and intra-cellular compartments. In contrast, Tryptophan concentration is upregulated in the cells and downregulated in the extracellular environment. (**Figure 3F**) Enrichment analysis indicated in correspondence of these metabolic signatures an alteration in *Glycine and Serine metabolism,* an imbalance of *Sphingolipid and phospholipid metabolism,* and most energetic pathways, such as the *Ketone body* and *Pyruvate metabolism, Citric acid,* and *Urea cycle.* (**Figure 3G**) ^44, 45^

### The benefits of nicotine in reducing Aβ (1-42) toxicity

As previously reported, a great deal of data has proved the benefits of nicotine in reducing Aβ (1-42) toxicity. Interestingly, in line with these data, our pharmacometabolomic study indicates a rebalance of the metabolic condition of SH-SY5Y cells pretreated with nicotine before being incubated with Aβ (1-42) towards the metabolic condition of the healthy control cells. (**Figure 4C, D**) A careful analysis of our data to understand how nicotine may impact different sides of cellular metabolism suggests significant effects of the alkaloid on (i) amino acid metabolism, particularly those involved with neurotransmission, (ii) energy metabolism (iii) membrane phospholipid metabolism.

### Nicotine treatment affects amino acid metabolism, particularly those involved in neurotransmission

Nicotine is effective in i) reducing Glutamate concentration in the extra- and intracellular compartments. Upregulation of excitotoxic glutamate transmission is a typical signature of neurodegenerative disease. ^46^ We demonstrated that the neuroprotective action of nicotine can be exerted through the down-regulation of glutamate pathways. ^47^ (**Figures 4C, D**, **E**) ii) increasing Serine concentration. Dietary supplementation with L-serine has been shown to restore both synaptic plasticity and memory alteration, suggesting it to be a potential therapy for AD. ^48^ Whereas deficiency of L-serine and its downstream products is associated with severe neurological deficits. ^49^ (**Figure 4B**) iii) increasing Arginine excretion, consistent with a rebalancing of the *iv)* reducing intracellular aromatic amino acids -Phenylalanine and Tryptophan- thus reverting an AD pathology signature, consisting in abnormally high concentrations of aromatic amino acids in the brain. ^50^ (**Figures 4D, E**).

### Nicotine treatment affects energetic metabolism

Nicotine in SH-SY5Y cells treated with Aβ (1-42) induces reduced levels of Acetoacetate, increased concentrations of 3- Hydroxybutyrate, and rebalance of *Pyruvate metabolism* and TCA (**Figures 4B, D, E**). High Pyruvate concentrations observed in Aβ (1-42)-treated cells are consistent with a downregulation of TCA and hypoglycemia condition (**Figure 3B, F, G**), as observed in the brains of AD patients. ^44^ Accordingly, the intracellular concentration of Glucose was shown to be reduced by Aβ (1-42) treatment (**Figure 3D**). This hypoglycaemic state is reversed in the presence of nicotine thanks to a shift in energy metabolism towards *Ketone body metabolism*, as shown by the increased conversion of Acetoacetate into 3-hydroxybutyrate. Thus, nicotine may protect cells from hypoglycemia by favoring the use of ketone bodies as an alternative fuel source. (**Figure 4E**) ^51^

Elevated homocysteine levels have been recognized as a contributing factor to the development of cognitive decline, dementia, and AD in elderly individuals since they induce mitochondria dysfunction. ^52^ Nicotine pre-treatment reduces Homocysteine and Betaine intracellular concentrations and increases Glutathione levels (**Figures 4B, D**). Indeed, a decrease in Betaine induces upregulation of Glutathione production, which modulates Homocysteine concentrations ^53^. Therefore, nicotine affects the *Folate pathway, Betaine metabolism, Homocysteine degradation,* and *Glutathione metabolism* (**Figure 4E**); all these effects are compatible with a final nicotine antioxidant effect.

Furthermore, nicotine restores Riboflavin physiological concentrations (**Figure 4B**). Because of riboflavin deficiency found in AD patients, therapies based on flavin mononucleotide (FMN) supplementation have an effect in containing AD symptoms. ^54^ Our results confirm the low riboflavin concentrations in the Aβ (1-42) pretreated cell model and show the efficacy of nicotine in rebalancing FMN physiological concentrations (**Figures 3F, 4B**).

### Nicotine treatment affects membrane phospholipid metabolism

Nicotine pre-treatment significantly upregulates *Phosphoethanolamine biosynthesis* while decreasing Glycerophosphocholine, a degradative product of phosphatidylcholine (**Figures 4B,E**) ^55^. Moreover, nicotine treatment increases Citicoline concentration (**Figure 4B**), an essential intermediate in the biosynthetic pathway of phosphatidylcholine in cell membranes. ^56^ Several studies have reported a dysregulation of phospholipid metabolism in AD, particularly an upregulation of phosphatidylcholine and phosphoethanolamine production ^57^. We demonstrate that nicotine restores phospholipid metabolism, which is significantly impaired in AD. (**Figure 4E**)

In conclusion, as shown in **Figure 5**, nicotine pre-treatment of SH-SY5Y cells exposed to Aβ (1-42) peptide induces i) glutamate reduction where excessive glutamate can lead to excitotoxicity, neuronal damage, and disease progression; ii) serine increasing, where low serine levels are linked to neurological disorders, cognitive decline or mood disorders; iii) improvement of mitochondrial function, counteracting hyperhomocysteinemia associated with neurodegeneration and fighting hypoglycemia with the ketone body consumption shift; iv) rebalance in membrane phospholipid metabolism, restoring the synthesis of phospho- and sphingolipids necessary for membrane integrity.

**Figure 5.**
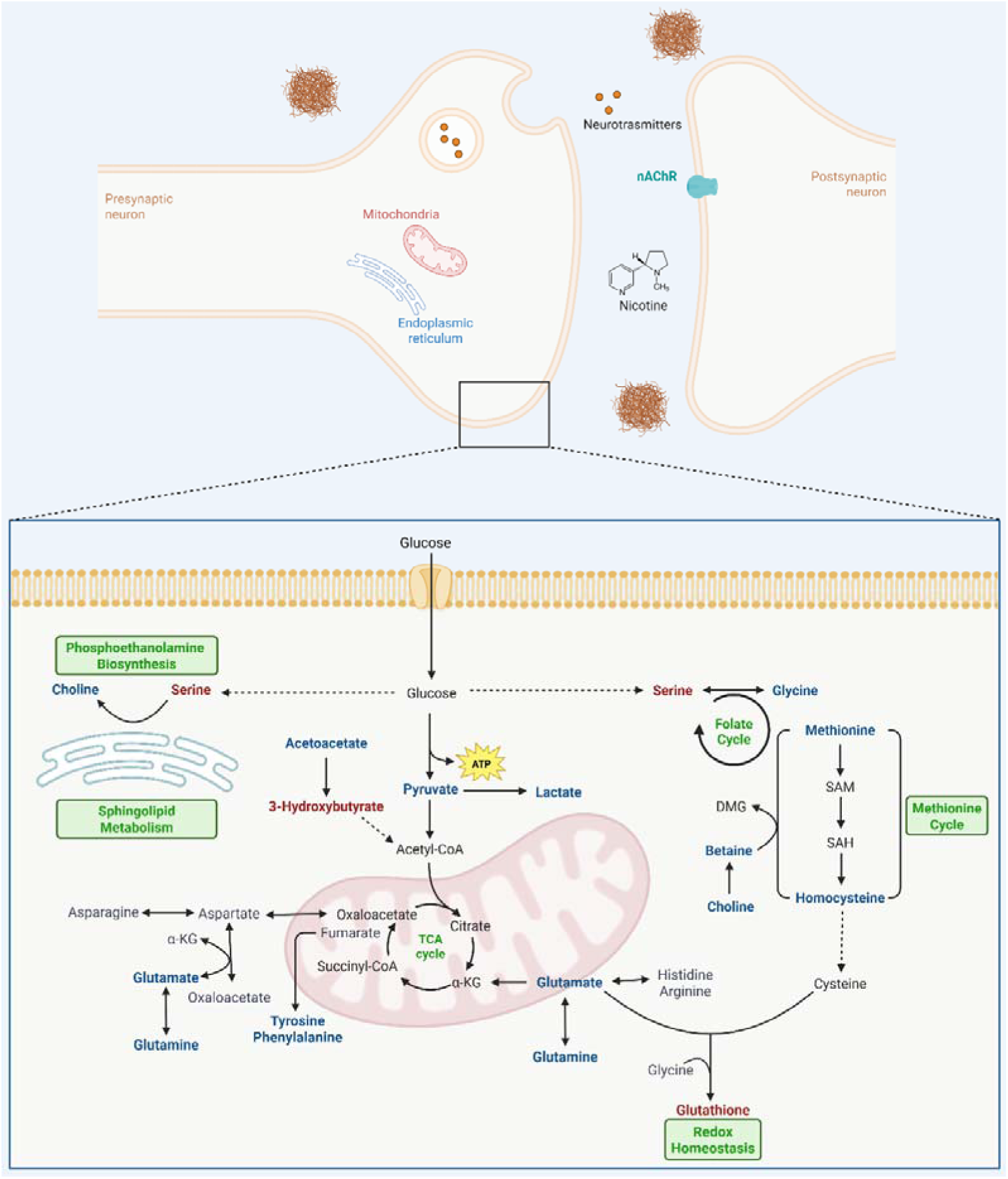
Overview of the metabolic effects of Nicotine on Alzheimer’s disease Upregulated metabolites in Aβ (1-42)-Nicotine cells are indicated in red; downregulated ones in blue; dysregulated pathways are labelled in green.

## Author Contributions

Conceptualization, A.M.D.; methodology, E,N., C.M., M.G., M.B; software, C.M. and E.N.; validation, E.N., C.M., M.G., M.B.; formal analysis, E.N., C.M., M.G.; investigation, E.N., C.M.; resources, A.M.D.; data curation, E.N.; writing—original draft preparation, E.N., C.M. and A.M.D.; writing—review and editing, E.N, C.M. and A.M.D.; visualization, E.N., A.M.D.; supervision, A.M.D.; project administration, A.M.D.; funding acquisition, A.M.D. All authors have read and agreed to the published version of the manuscript.

## SUPPORTING INFORMATION

The following supporting information is available free of charge at ACS we site http://pubs.acs.org

**Figure S1.**
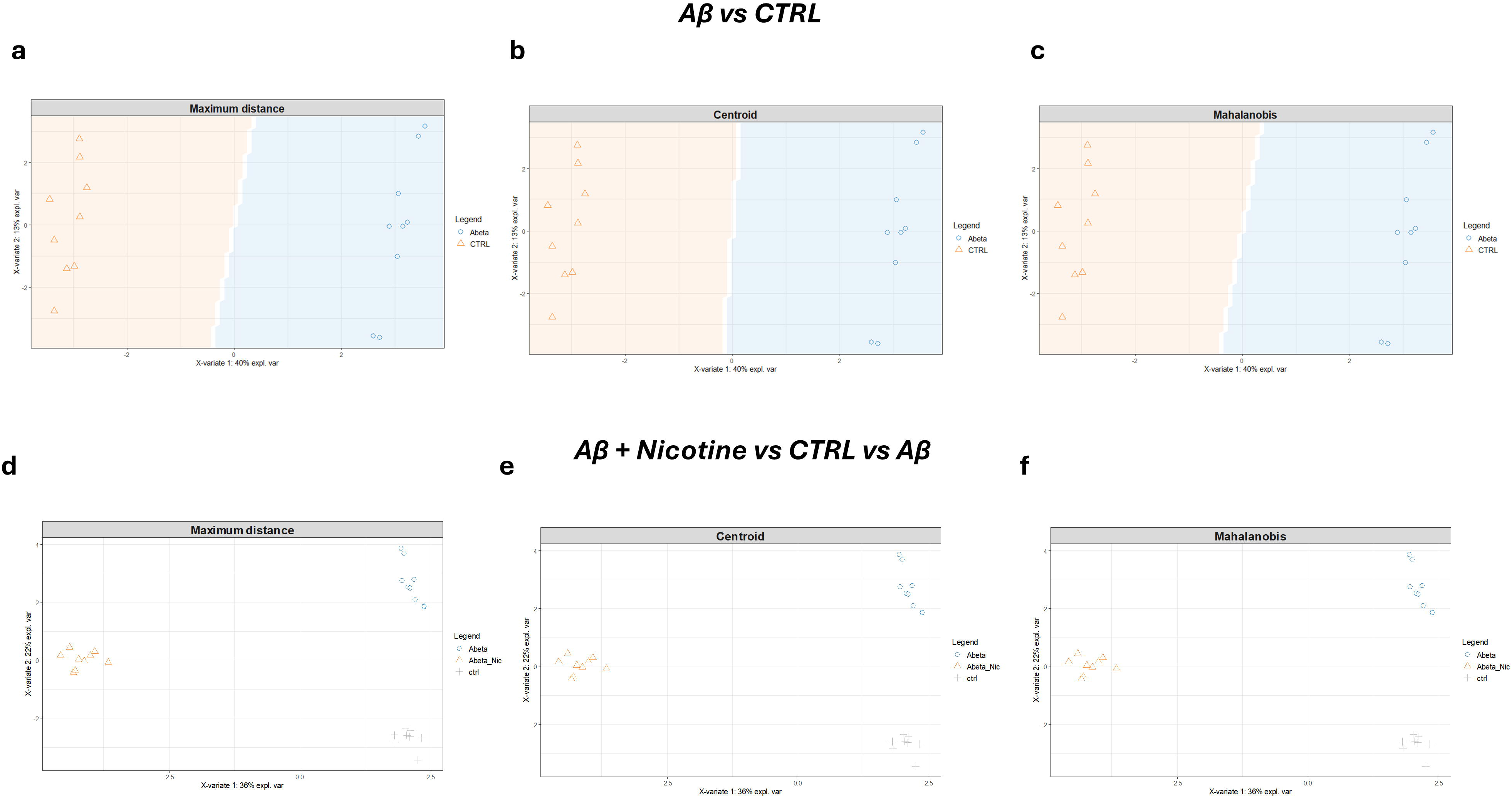
Sample prediction area plot created using Maximum distance, Centroid and Mahalanobis showing the distribution of samples in validation areas related to Aβvs CTRL (**a,b,c**) and Aβ+Nicotine vs CTRL vs Aβ (**e,f,g**).

**Figure S2.**
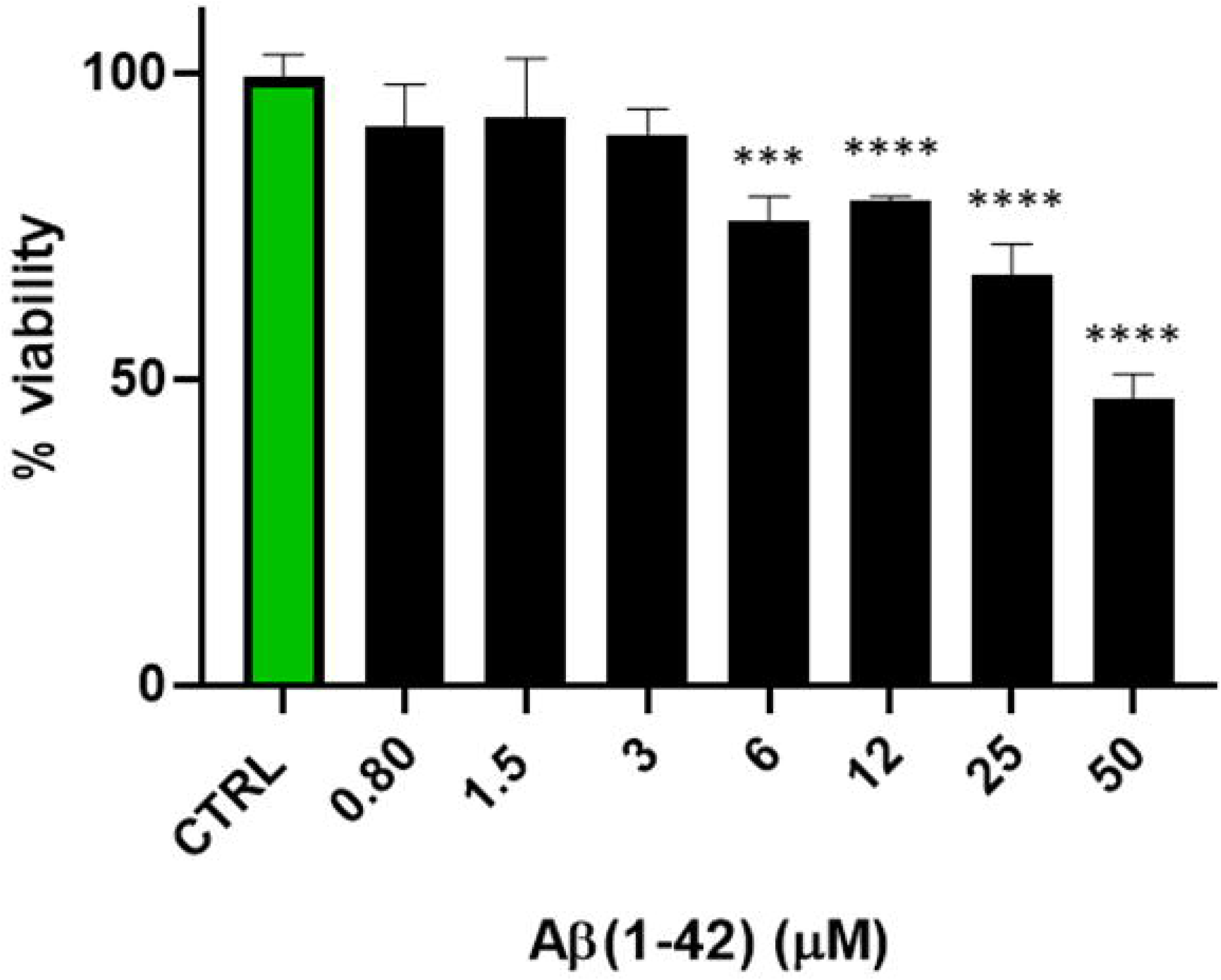
Aβ (1-42) effect on SH-SY5Y cells viability after 48h, examined by the CCK-8 assay. The viability variations were calculated as the percentage of viable cells in treated cultures compared to untreated ones (CTRL). Results are shown as mean ± standard deviation (SD) from three independent experiments. ***, **** denote respectively p < 0.001 and p < 0.0001 vs CTRL.

**Table S1**. Pathway Enrichment analysis discriminates between the analysed clusters. The number of hits corresponds to the number of metabolites detected in the spectrum that participate in the biochemical pathways and are explicit in the column ‘metabolites’. Raw p represents the significance validation index reporting the p-value; Holm Bonferroni represents the adjustment of the p.value for the number of analysed samples (Holm p.); the FDR index calculates the number of False Discovery Rates. Biochemical pathways with hits>2 and Raw.p, Holm p, FDR <0.05 were considered significant.

## ABBREVIATIONS

AD: Alzheimer’s disease
Aβ: Amyloid peptide
nAChRs: ACh receptors
MAPKs: MAP kinases
TSP-d_4_: trimethylsilyl propanoic-2,2,3,3-d_4_ acid
MVA: Multivariate analysis
s-PLS-DA: Sparse Partial Least Squares
PCA: Principal Component Analysis
CCA: Canonical correspondence analysis
NAA: N-Acetylaspartate

## Data availability

Metabolomics data have been deposited to the EMBL-EBI MetaboLights database (https://www.ebi.ac.uk/metabolights/) with the identifier MTBLS12706.

## For TOC Only

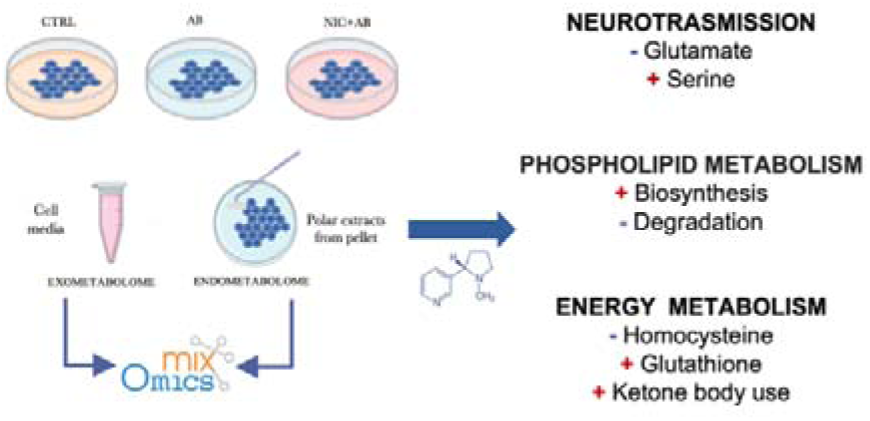

