## Supplementary material for "Analyzing nicotine action against amyloid toxicity by NMR-pharmacometabolomics: an exploratory study": Figures S1, S2 and Table S1

**Table of contents**

**Figure S1**. Sample prediction area plot created using Maximum distance, Centroid and Mahalanobis showing the distribution of samples in validation areas related to Aβvs CTRL (**a,b,c**) and Aβ+Nicotine vs CTRL vs Aβ (**e,f,g**).

**Figure S2.** Aβ (1-42) effect on SH-SY5Y cells viability after 48h, examined by the CCK-8 assay. The viability variations were calculated as the percentage of viable cells in treated cultures compared to untreated ones (CTRL). Results are shown as mean ± standard deviation (SD) from three independent experiments. ***, **** denote respectively p < 0.001 and p < 0.0001 vs CTRL.

**Table S1.** Pathway Enrichment analysis discriminates between the analysed clusters. The number of hits corresponds to the number of metabolites detected in the spectrum that participate in the biochemical pathways and are explicit in the column ‘metabolites’. Raw p represents the significance validation index reporting the p-value; Holm Bonferroni represents the adjustment of the p.value for the number of analysed samples (Holm p.); the FDR index calculates the number of False Discovery Rates. Biochemical pathways with hits>2 and Raw.p, Holm p, FDR <0.05 were considered significant.

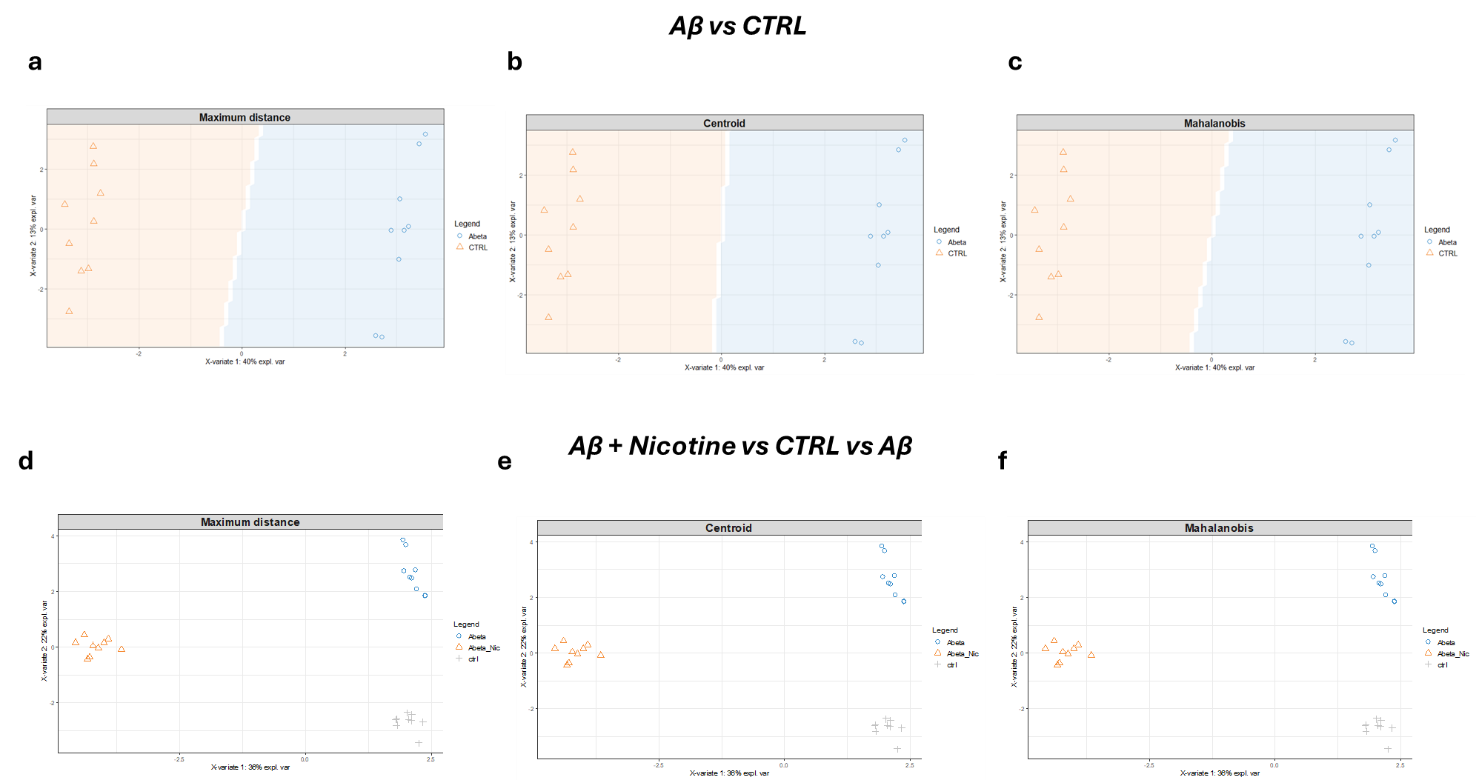

**Figure S1**. Sample prediction area plot created using Maximum distance, Centroid and Mahalanobis showing the distribution of samples in validation areas related to Aβ vs CTRL (**a,b,c**) and Aβ+Nicotine vs CTRL vs Aβ (**e,f,g**).

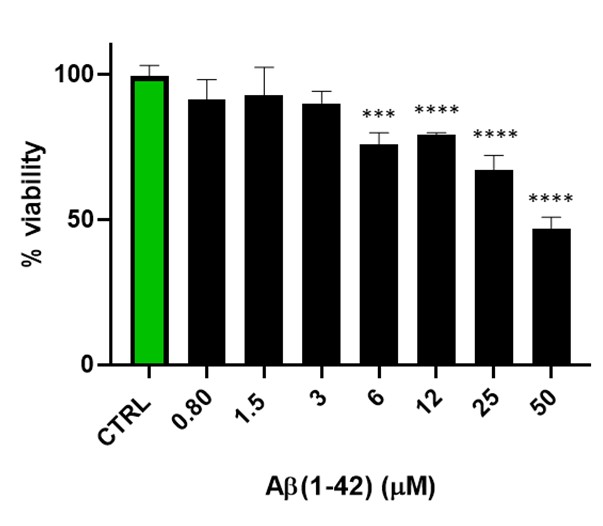

**Figure S2.** Aβ (1-42) effect on SH-SY5Y cells viability after 48h, examined by the CCK-8 assay. The viability variations were calculated as the percentage of viable cells in treated cultures compared to untreated ones (CTRL). Results are shown as mean ± standard deviation (SD) from three independent experiments. ***, **** denote respectively p < 0.001 and p < 0.0001 vs CTRL

**Table S1.** Pathway Enrichment analysis discriminates between the analysed clusters. The number of hits corresponds to the number of metabolites detected in the spectrum that participate in the biochemical pathways and are explicit in the column ‘metabolites’. Raw p represents the significance validation index reporting the p-value; Holm Bonferroni represents the adjustment of the p.value for the number of analysed samples (Holm p.); the FDR index calculates the number of False Discovery Rates. Biochemical pathways with hits>2 and Raw.p, Holm p, FDR <0.05 were considered significant.

| **Pathways Aβ vs CTRL** | **Hits** | **Raw p** | **Holm p** | **FDR** | **Metabolites** |
| --- | --- | --- | --- | --- | --- |
| Phosphatidylethanolamine Biosynthesis | 3 | 4,58E-06 | 3,94E-04 | 3,94E-04 | Choline Serine ATP |
| Citric Acid Cycle | 3 | 1,23E-05 | 9,90E-03 | 1,32E-03 | Pyruvic acid; Succinic acid; Adenosine triphosphate |
| Glutamate Metabolism | 8 | 1,36E-05 | 1,16E-03 | 5,85E-04 | Glycine,Glutathione; Glutamic acid,L-Aspartic acid,Pyruvic acid;Succinic acid;Adenosine triphosphate;Glutamine |
| Ketone Body Metabolism | 3 | 2,78E-05 | 2,33E-03 | 6,36E-05 | 3-Hydroxybutyric acid; Acetoacetic acid; Succinic acid; |
| Glycine and Serine Metabolism | 12 | 2,96E-05 | 2,45E-03 | 6,36E-05 | 3-Hydroxybutyric acid; Acetoacetic acid; Succinic acid |
| Warburg Effect | 7 | 6,06E-05 | 4,97E-03 | 1,04E-03 | 2-Ketobutyric acid; Betaine;Glycine;Glutamic acid;L-Threonine; Serine;Pyruvic acid;Sarcosine;L-Arginine; Adenosine triphosphate;Methionine; Homocysteine |
| Mitochondrial Electron Transport Chain | 2 | 1,22E-04 | 9,90E-03 | 1,32E-03 | Succinic acid; Adenosine triphosphate |
| Phytanic Acid Peroxisomal Oxidation | 2 | 1,22E-04 | 9,90E-03 | 1,32E-03 | Succinic acid; Adenosine triphosphate |
| Butyrate Metabolism | 3 | 1,10E-03 | 8,56E-02 | 1,05E-02 | Acetoacetic acid;  Succinic acid; Adenosine triphosphate |
| Methionine Metabolism | 9 | 1,64E-03 | 1,26E-01 | 1,41E-02 | 2-Ketobutyric acid; Betaine; Choline; Glycine; Serine;  Sarcosine; Adenosine triphosphate; Methionine; Homocysteine |
| Sphingolipid Metabolism | 4 | 2,03E-03 | 1,54E-01 | 1,59E-02 | D-Glucose;Serine; Uridine diphosphate glucose;Adenosine triphosphate |
| Carnitine Synthesis | 4 | 2,48E-03 | 1,86E-02 | 1,78E-02 | L-Carnitine; Glycine; Lysine; Succinic acid |
| Ammonia Recycling | 8 | 3,08E-03 | 2,28E-02 | 2,04E-02 | Glycine; Glutamic acid; Histidine; Serine; L-Aspartic acid;Pyruvic acid; Adenosine triphosphate; Glutamine; |
| Oxidation of Branched Chain Fatty Acids | 3 | 3,98E-03 | 2,91E-02 | 2,45E-02 | L-Carnitine;Succinic acid; Adenosine triphosphate |
| Glucose-Alanine Cycle | 3 | 7,58E-03 | 5,46E-02 | 4,35E-02 | D-Glucose; Glutamic acid; Pyruvic acid |
| Cysteine Metabolism | 3 | 8,51E-03 | 6,04E-03 | 4,58E-02 | D-Glucose; Glutamic acid; Pyruvic acid |
| Alanine Metabolism | 4 | 1,07E-02 | 7,49E-03 | 5,39E-02 | Glycine; Glutamic acid;Pyruvic acid;Adenosine triphosphate |
| Pyruvaldehyde Degradation | 2 | 1,13E-02 | 7,79E-03 | 5,39E-02 | Glutathione; Pyruvic acid |
| Pyruvate Metabolism | 5 | 1,61E-02 | 0.00010966 | 7,30E-02 | Acetic acid; Glutathione; ; Lactic acid; Pyruvic acid; Adenosine triphosphate |
| Glycolysis | 3 | 1,90E-02 | 0.00012722 | 7,78E-02 | D-Glucose;Pyruvic acid;  Adenosine triphosphate; |
| Transfer of Acetyl Groups into Mitochondria | 3 | 1,90E-02 | 0.00012722 | 7,78E-02 | D-Glucose; Pyruvic acid; Adenosine triphosphate; |
| Urea Cycle | 6 | 2,34E-02 | 0.0001519 | 8,83E-02 | Glutamic acid; L-Aspartic acid; Pyruvic acid; L-Arginine; Adenosine triphosphate;  Glutamine |
| Gluconeogenesis | 4 | 2,36E-02 | 0.0001519 | 8,83E-02 | D-Glucose;  Lactic acid;  Pyruvic acid;  Adenosine triphosphate |
| **Pathways**  **Aβ+Nicotine**  **vs CTRL vs Aβ** | **Hits** | **Raw p** | **Holm p** | **FDR** | **Metabolites** |
| Amino Sugar Metabolism | 7 | 3,07E-14 | 2,64E-12 | 1,44E-12 | Acetic acid; Glutamic acid; Pyruvic acid; Pyrophosphate; Adenosine triphosphate; Glutamine; D-Fructose |
| Glutamate Metabolism | 8 | 3,62E-14 | 3,08E-12 | 1,44E-12 | Glycine; Glutathione; Glutamic acid; Pyruvic acid; Succinic acid; Phosphoribosyl pyrophosphate; Adenosine triphosphate; Glutamine |
| Warburg Effect | 7 | 5,04E-14 | 4,23E-12 | 1,44E-12 | D-Glucose; Glutamic acid;Lactic acid; Pyruvic acid; Succinic acid; Adenosine triphosphate;Glutamine; |
| Citric Acid Cycle | 3 | 7,66E-14 | 6,36E-12 | 1,65E-12 | Pyruvic acid; Succinic acid; Adenosine triphosphate; |
| Ketone Body Metabolism | 3 | 7,11E-13 | 5,83E-11 | 1,22E-11 | 3-Hydroxybutyric acid; Acetoacetic acid; Succinic acid; |
| Arginine and Proline Metabolism | 7 | 9,29E-13 | 7,53E-11 | 1,33E-11 | Glycine; Glutamic acid; Proline; L-Aspartic acid; Succinic acid; L-Arginine; Adenosine triphosphate |
| Ammonia Recycling | 8 | 1,91E-11 | 1,52E-09 | 2,31E-10 | Glycine; Glutamic acid; Histidine; Serine; L-Aspartic acid; Pyruvic acid; Adenosine triphosphate; Glutamine |
| Butyrate Metabolism | 3 | 2,15E-11 | 1,69E-09 | 2,31E-10 | Acetoacetic acid; Succinic acid; Adenosine triphosphate; |
| Betaine Metabolism | 5 | 4,19E-11 | 3,27E-09 | 4,00E-10 | Betaine; Choline; Adenosine triphosphate; Methionine; Homocysteine; |
| Glycine and Serine Metabolism | 12 | 9,22E-11 | 7,10E-10 | 7,93E-10 | 2-Ketobutyric acid; Betaine; Glycine; Glutamic acid; Pyruvic acid; Sarcosine; L-Arginine; Adenosine triphosphate; Methionine; Homocysteine; |
| Urea Cycle | 6 | 1,11E-10 | 8,47E-09 | 8,72E-10 | Glutamic acid;L-Aspartic acid;Pyruvic acid; L-Arginine; Adenosine triphosphate; Glutamine; |
| Carnitine Synthesis | 4 | 2,08E-10 | 1,56E-08 | 1,49E-09 | L-Carnitine; Glycine; Lysine; Succinic acid; |
| Oxidation of Branched Chain Fatty Acids | 3 | 2,58E-11 | 1,91E-08 | 1,71E-09 | L-Carnitine; Succinic acid; Adenosine triphosphate; |
| Glycolysis | 3 | 8,64E-10 | 6,22E-08 | 4,64E-09 | D-Glucose; Pyruvic acid; Adenosine triphosphate |
| Transfer of Acetyl Groups into Mitochondria | 3 | 8,64E-10 | 6,22E-08 | 4,64E-09 | D-Glucose; Pyruvic acid; Adenosine triphosphate; |
| Gluconeogenesis | 4 | 1,96E-09 | 1.33e-11 | 8,85E-09 | D-Glucose; Lactic acid; Pyruvic acid; Adenosine triphosphate; |
| Cysteine Metabolism | 3 | 3,96E-09 | 2,57E-07 | 1,55E-08 | Glutamic acid; Pyruvic acid; Adenosine triphosphate |
| Glucose-Alanine Cycle | 3 | 6,44E-09 | 4,12E-07 | 2,31E-08 | D-Glucose; Glutamic acid; Pyruvic acid; |
| Alanine Metabolism | 4 | 6,45E-09 | 4,12E-07 | 2,31E-08 | Glycine; Glutamic acid; L-Alanine; Pyruvic acid; Adenosine triphosphate |
| Pyruvate Metabolism | 5 | 1,21E-08 | 7,41E-07 | 4,02E-08 | Acetic acid; Glutathione;Lactic acid; Pyruvic acid; Adenosine triphosphate |
| Aspartate Metabolism | 7 | 1,81E-08 | 1,09E-06 | 5,77E-08 | Acetic acid, Glutamic acid; L-Aspartic acid; L-Arginine; Adenosine triphosphate; Glutamine; N-Acetyl-L-aspartic acid |
| Nicotinate and Nicotinamide Metabolism | 3 | 4,17E-08 | 2,46E-06 | 1,28E-07 | Glutamic acid; Adenosine triphosphate; Glutamine; |
| Valine, Leucine and Isoleucine Degradation | 8 | 6,10E-08 | 3,54E-06 | 1,81E-07 | Acetoacetic acid; Glutamic acid; Isoleucine; Methylmalonic acid; Succinic acid;Adenosine triphosphate; Leucine; L-Valine; |
| Tryptophan Metabolism | 4 | 7,06E-08 | 3,95E-06 | 1,96E-08 | Formic acid; Glutamic acid; Adenosine triphosphate; L-Tryptophan |
| Methionine Metabolism | 9 | 7,93E-08 | 4,36E-06 | 2,13E-07 | 2-Ketobutyric acid; Betaine; Choline; Glycine; Serine; Sarcosine; Adenosine triphosphate; Methionine; Homocysteine; |
| Purine Metabolism | 5 | 1,06E-07 | 5,74E-06 | 2,77E-07 | Glycine; Glutamic acid; L-Aspartic acid; Adenosine triphosphate; Glutamine |
| Homocysteine Degradation | 3 | 1,40E-06 | 7,12E-05 | 3,34E-06 | 2-Ketobutyric acid; Serine; Homocysteine |
| Phosphatidylethanolamine Biosynthesis | 3 | 2,20E-06 | 1,01E-04 | 4,62E-06 | Choline; Serine; Adenosine triphosphate; |
| Tyrosine Metabolism | 4 | 2,79E-06 | 1,23E-04 | 5,58E-06 | Acetoacetic acid; Glutamic acid; L-Tyrosine; L-Aspartic acid; |
| Fatty Acid Biosynthesis | 3 | 5,72E-06 | 2,46E-04 | 1,12E-05 | Acetic acid; Acetoacetic acid; 3-Hydroxybutyric acid; |
| Sphingolipid Metabolism | 4 | 7,82E-06 | 3,28E-04 | 1,47E-05 | D-Glucose; Serine;Uridine diphosphate glucose; Adenosine triphosphate; |
| Glutathione Metabolism | 5 | 8,50E-06 | 3,40E-04 | 1,56E-05 | Glycine; Glutathione; Glutamic acid; Pyroglutamic acid; Adenosine triphosphate |
| Folate Metabolism | 3 | 1,20E-05 | 4,69E-04 | 2,16E-05 | Formic acid; Glutamic acid; Adenosine triphosphate |
| Selenoamino Acid Metabolism | 3 | 1,75E-05 | 6,65E-04 | 3,07E-05 | 2-Ketobutyric acid; Serine; Adenosine triphosphate |
| Phenylalanine and Tyrosine Metabolism | 5 | 3,01E-05 | 1,11E-03 | 5,17E-05 | Acetoacetic acid; Glutamic acid; L-Tyrosine; Phenylalanine; Adenosine triphosphate; |
| Propanoate Metabolism | 6 | 1,20E-04 | 4,32E-03 | 2,02E-04 | 2-Ketobutyric acid; 2-Hydroxybutyric acid; Glutamic acid; Methylmalonic acid; Adenosine triphosphate; L-Valine |
